# The calcifying interface in a stony coral’s primary polyp: An interplay between seawater and an extracellular calcifying space

**DOI:** 10.1101/2021.06.09.447817

**Authors:** Gal Mor Khalifa, Shani Levy, Tali Mass

## Abstract

Stony coral exoskeletons build the foundation to the most biologically diverse yet fragile marine ecosystems on earth, coral reefs. Understanding corals biomineralization mechanisms is therefore crucial for coral reef management and for using coral skeletons in geochemical studies. In this study, we combine in-vivo and cryo-electron microscopy with single-cell RNA-seq data to gain novel insights into the calcifying micro-environment that facilitates biomineralization in primary polyps of the stony coral *Stylophora pistillata*. We show an intimate involvement of seawater in this micro-environment. We further document increased tissue permeability and a highly dispersed cell packing in the tissue secreting the coral skeleton (i.e. calicoblastic). We also observe an extensive filopodial network containing carbon-rich vesicles extruding from some of the calicoblastic cells. Single-cell RNA-Seq data interrogation shows that calicoblastic cells express genes involved in filopodia and vesicle structure and function. These observations provide a new conceptual framework for resolving stony corals biomineralization processes.

## Introduction

Coral reefs are highly biodiverse ecosystems (*1*–*3*). Stony corals inhabiting these ecosystems produce calcium carbonate exoskeletons which, on geological time scales, can lead to the formation of massive coral reefs spanning thousands kilometers in shallow tropical and subtropical seas (*4, 5*). Coral reef ecosystems around the world are facing major threats due to multiple global anthropogenic stressors including sea surface warming and ocean acidification, and local anthropogenic stressors such as overfishing, pollution, marine construction and diving pressure (*6*–*11*). Coral reef risk assessment is a highly complex task, not only because of the multiple stressors involved, but also due to the lack of mechanistic understanding of some physiological processes in corals, including how they build their exoskeletons. Coral skeletogenesis is a biologically controlled process performed by the coral animal. Therefore, corals can respond to and compensate for environmental changes such as ocean acidification to some extent (*12*). It is also reported that some coral reef ecosystems are more resistant than others to episodic sea surface warming (*13*). A major missing link for predicting the degree of stony coral resilience to environmental changes is the understanding of the basic mechanisms, at the tissue, cellular and molecular levels, by which corals calcify. Stony coral skeletons are made of calcium carbonate almost entirely of the polymorph aragonite (*14, 15*). The elemental and isotopic compositions of the aragonitic coral skeletons record the external seawater chemistry in which they were formed, but with an offset due to the biological control of the coral animal over this biomineralization process. This offset is consistent within individual species and termed the vital effect e.g. (*16*). Coral skeletons are therefore, used for reconstructing recent and past ocean chemistry and climate (*16*). Resolving the biomineralization process in stony corals is therefore, essential for better understanding the use of coral skeleton elemental and isotopic composition for ocean chemistry reconstructions, as well as for assessing coral reef fate under current and future climate conditions (*17*).

Most scleractinian or stony corals (Anthozoa) form colonies with the basic unit of a polyp. A coral’s life cycle involves a swimming planula which metamorphoses, settles and immediately commences rapid calcification in order to attach firmly to the substrate and to form the primary polyp (*18*). Polyps have a cylindrical shape with a central mouth surrounded by tentacles used for predation (Fig 1a, e). Many stony corals also contain symbiotic dinoflagellates of the family Symbiodiniaceae (*19*) and feed both by predation and on photosynthetic products supplied by their endosymbionts (*20*). Coral anatomy includes two body layers, an ectoderm and an endoderm, separated with a non-cellular gelatinous layer called mesoglea. This anatomy repeats itself in the oral tissue that faces the external seawater, and in the aboral tissue that faces the exoskeleton (*4, 21, 22*). The oral ectoderm contains stinging cells (nematocytes) which play a role in predation and defense while the aboral ectoderm contains calicoblastic cells responsible for secreting the mineralized exoskeleton within the extra-cellular medium (ECM) (*4, 22*). The skeleton of an individual polyp (the cup-shaped portion of the skeleton called a corallite) is composed of radially aligned plates (septa) projecting upwards from the base (Fig 1a, b). Each septum micro-structure includes the first-formed center of calcification (CoC) characterized by a nano-granular texture, while the rest of the septum is composed of fibrous elongated micron-sized single crystals arranged in a three-dimensional fan around the CoC (*14, 23*–*25*). The calicoblastic cell layer, the ECM and the exoskeletal mineral surface together create the micro-environment in which coral biomineralization occurs. Detailed characterization of this micro-environment is essential for resolving the biomineralization strategies in stony corals. Morphological and spatial characterization of the calicoblastic cell layer and the ECM have been obtained over the past few decades largely using conventional scanning electron microscopy (SEM), transmission electron microscopy (TEM) thin sections and histology in several stony coral species (*22, 26, 27*). However, as with all biological samples, these tissues are predominately composed of water-based solutions, the characterization of which is problematic using the above techniques. The native state volume and chemical composition of water-based solutions cannot be accurately characterized using the above techniques because of multiple solution exchange steps involved in the preparation procedures in which the native water-based solutions are effectively removed from the sample and replaced with preparative agents. Furthermore, chemical fixation, demineralization, staining, dehydration, and plastic or paraffin embedding involved in the preparation procedures may result in further morphological alterations of the tissue (*28*). In contrast, advances in cryo-fixation and cryo-electron microscopy (cryo-EM) techniques in recent years allow for observation of fully hydrated biological samples without the need of chemical fixation or staining. Great strides have been made in morphologic and spectroscopic analysis of cryo-preserved biological specimens since the first low temperature imaging of coral tissue and skeleton fragments in 2002 (*28*). This includes the use of high-pressure freezing as a cryo-fixation technique, which keeps the water molecules of the sample in a non-crystalline amorphous solid state, i.e., ‘vitrified’. This is due to the very short duration of this freezing procedure (miliseconds) allowed by the high-pressure conditions, which leaves no time for ice crystals to form within the sample (at least not to a size larger than a few nm, which is the spatial resolution of cryo-SEM imaging). Ice crystals formed during longer freezing procedures can change both the structure and local chemical composition of the sample. Therefore a high-pressure frozen sample that is imaged using cryo-SEM more closely represents in-vivo conditions than alternative techniques mentioned above (*29, 30*). Cryo-EM also allows both a large imaged field of view and high-resolution subcellular morphological characterization of the sample (*31, 32*). When coupled with cryo-planing this preparation technique further provides a sample surface ideal for cryo-elemental analysis (cryo-energy dispersive x-ray spectroscopy-cryo-EDS). Cryo-SEM/EDS is one of only a few techniques available today that allows in-situ elemental analysis of soft tissues and their water-based solution components (*33, 34*) in addition to the widely characterized elemental composition of the hard mineralized coral exoskeleton (*35, 36*). We therefore adopted cryo-SEM/EDS analysis to characterize the calcifying tissue-mineral interface in this study.

**Fig. 1:**
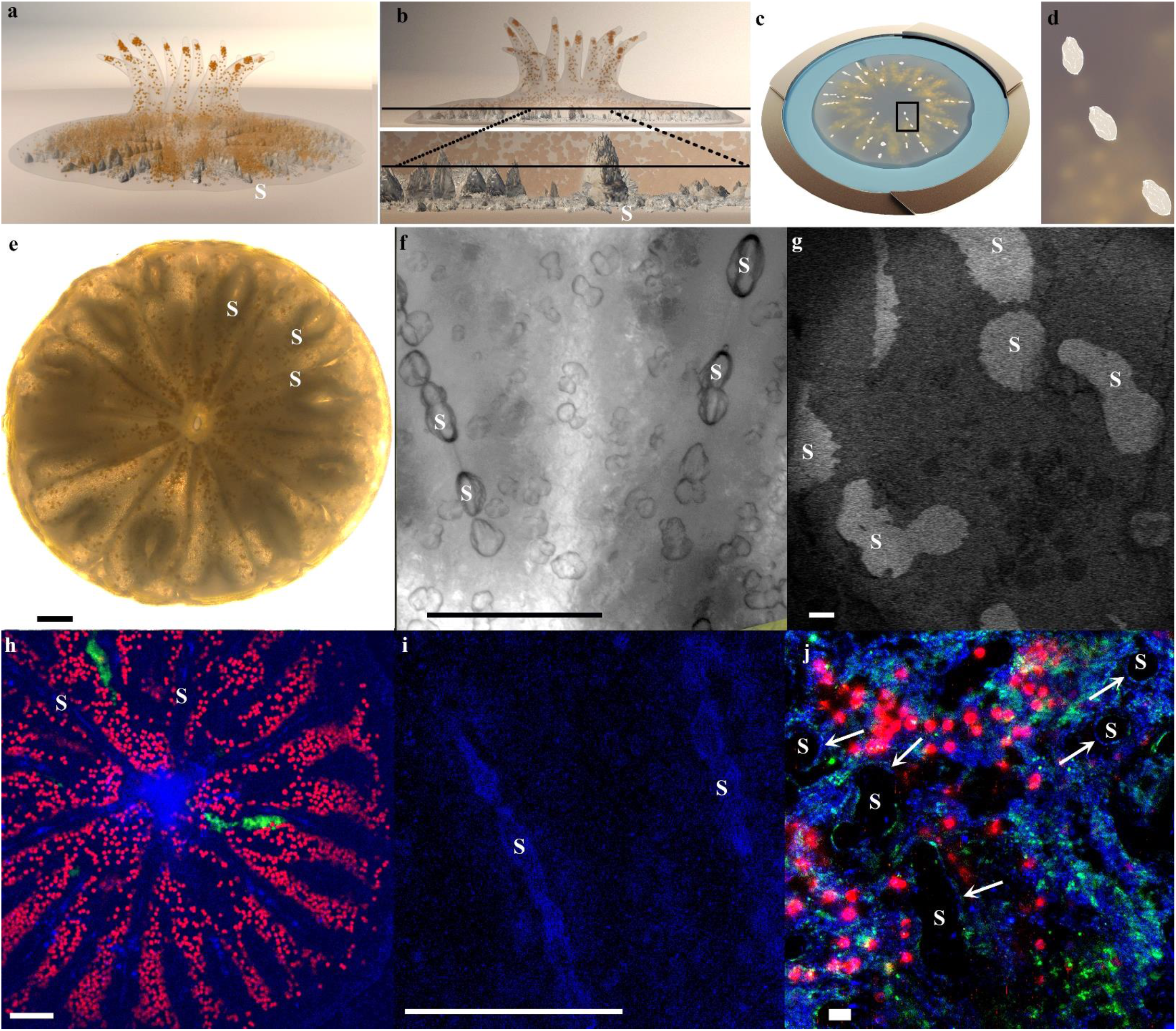
A Primary polyp and its forming mineralized septa. (a) Illustration of a few days old primary polyp with 12 tentacles. Tissue is colored in transparent grey, endosymbionts in orange and mineral in white. (b) Upper panel: An illustration of the polyp (side view) internal plane (black line) revealed by cryo-planing. Bottom panel: A magnification of this same plane along two of the polyp septa. (c) Illustration of the high-pressure frozen, cryo-planed primary polyp (top view) vitrified in natural seawater (blue) inside a high-pressure freezing disc (gold). (d) Magnification of the area marked with black rectangle in (c). (e) Light microscopy image of five days old primary polyp. (f) Higher magnification wide field microscope image of two forming septa in the primary polyp. (g) Cryo-SEM (ESB mode) micrograph of a high-pressure frozen freeze fractured primary polyp showing a non-continuous fracture surface of the forming septa. Septa mineral surface appears white and coral tissue surface appears grey. (h) Confocal laser scanning microscope overview image of a calcein-blue labeled primary polyp. Area of forming septa and mouth cavity are labeled in blue (calcein). Coral tissue auto-fluorescence is green and photosymbionts auto-fluorescence is red. (i) Higher magnification of two forming septa labeled with calcein blue (blue channel). (j) Cryo-fluorescence image of the same freeze fractured primary polyp as in (g) imaged in blue, green and red channels. Thin calcein blue labeled layer limning of the external surface of some septa is depicted with white arrows. **S**-septum mineral, **T**- tentacle. Scale bars: e, f, h, i- 100 µm, g, j- 20 µm.

Another cutting-edge technique recently applied for the first time in stony corals is single cell RNA sequencing (scRNA-seq) (*37*), which reveals cellular specialization in stony corals. The new *S. pistillata* cell atlas shows the transcriptional profile of the calicoblastic cells, among other cell types, in the primary polyp (*37*) and thus, provides insights into the molecular basis of the biomineralization processes carried out by these cells. Over 800 genes were found to be specific to the calicoblastic cells including those associated with biomineralization. Such as, carbonic anhydrase facilitating the inter-conversion of CO_2_to HCO_3-_ and HCO_3-_transporters (Fig. S1 (*37*)), and acid rich proteins found in the coral skeletal proteomes (*38*– *41*). The recent scRNA-seq analysis of multiple *S. pistillata* life stages also showed that 30% of the primary polyp cells are calicoblasts, while adult colonies are comprised of less than 7% calicoblasts cells (*37*). In addition, the authors reported an increased expression of biomineralization related genes in primary polyps compared with adults (Fig. S1 and (*37*)), likely supporting the assumption that mineralization activity is more rapid in the primary polyp stage compared the adult life stage (*18, 21*). This makes primary polyps useful targets of study to characterize the mineralization micro-environment and are therefore the subjects of this study.

Primary polyps (Fig 1. a, e) were high-pressure frozen and the mineral and its adjacent tissue were exposed for cryo-SEM/EDS analysis using either freeze-fracture to obtain topographic representation of the tissue (*32*), or cryo-planing (Fig. 1 b,c and d) to reveals the internal content of cells and intracellular structures for both imaging (cryo-SEM) and quantitative elemental analysis (cryo-EDS) (*33, 34*). We combine these observations with in-vivo dynamic fluorescence imaging and further used and re-analyzed the recently published calicoblasts transcriptional profile (*37*) as complementary data to our structural and elemental analyses. Our combination of two cutting-edge techniques, cryo-SEM/EDS and scRNA-seq analysis and in-vivo imaging, allows for the novel description of the micro-environment in which coral biomineralization occurs and the cells that execute this process. It also provides new insight into the mechanism of biomineralization and a conceptual framework to resolve the involvement of external seawater in this process. The latter is key for understanding the use of coral skeletons for reconstructing past ocean chemistry as well as for understanding the impact of current changes in ocean chemistry on the survival of stony corals as individuals and of coral reefs as a whole.

## Results

### Morphological characterization of primary polyp tissues using cryo-SEM

Primary polyps were studied 4-5 days after settlement. At this life stage, the primary polyp is adhered to the substrate, tentacles have formed and can be extended by the animal (Fig. 1a) and skeletogenesis is at its early stages in the sessile organism. The basal part of the primary polyp skeleton, termed the ‘basal plate’, is not yet formed (Fig 1a), and septa are often not fully developed, appearing non-continuous by conventional light microscopy (Fig. 1 e,f). A cryo-SEM energy-selective back scattered electron (ESB) mode image of a high-pressure-frozen, freeze-fractured primary polyp reveals the septum with non-continuous mineral surfaces spread along its long axis (Fig. 1g). Mineral surfaces are detectable with higher contrast (white) than adjacent tissue (grey) using the ESB detector that is sensitive to electron density. We used the fluorescent probe calcein to confirm that studied primary polyps were indeed depositing new mineral to thicken their septa at the time of cryo-fixation. Calcein is a cell impermeable dye (as oppose to the permeable version of this dye, calcein-AM) which when applied in the seawater is incorporated into the newly formed mineral and labels it (*42*). Primary polyps were labeled in-vivo with calcein blue and imaged both in-vivo (Fig. 1 h, i) and after cryo-fixation and freeze fracturing by cryo-fluorescence using a confocal laser scanning microscope combined with a cryo-stage (Fig. 1j). In both in-vivo and cryo-fluorescence images a thin calcein blue layer is observed lining the external septa surfaces, indicative of newly formed mineral which was deposited during the labeling period. In the freeze-fractured cryo-fluorescence image, the bulk internal part of the fractured mineral surfaces appears black due to the absence of any fluorescence labeling inside the skeleton while newly formed calcein labeled layer is observed only at the periphery of the fractured surface.

Studying the freeze-fractured surface of primary polyps, we observed characteristic cells of the oral tissue such as nematocysts, mucus forming cells (mucocytes) and host cells containing symbiotic algae (Supplementary Fig. S1) (*22, 43*). We further observe the traditionally classified two body layers (*22*) of the aboral tissue adjacent with the mineral (Fig 2a). The mineral is indicated by its micro-crystalline structure (Fig. 2b) whereas both aboral body layers are indicated by their cellular composition in which nuclei and cellular membranes are clearly detected (Fig. 2c), and the non-cellular mesoglea separating them is identified by its rough surface texture (Fig 2d) characteristic of a vitrified solution (Appendix A figure S5 in *33*). It is noteworthy that not all observed areas are characterized with this classical tissue arrangement. Similar to the reduction of tissue layers near a skeletal spine observed by Tambutte et al. (*26*) in which the authors found only two ectodermal layers separated by mesoglea with no endodermal layers, in some of our observed loci nematocytes were found in close proximity with the mineral (Supplementary Fig. S2).

**Fig. 2:**
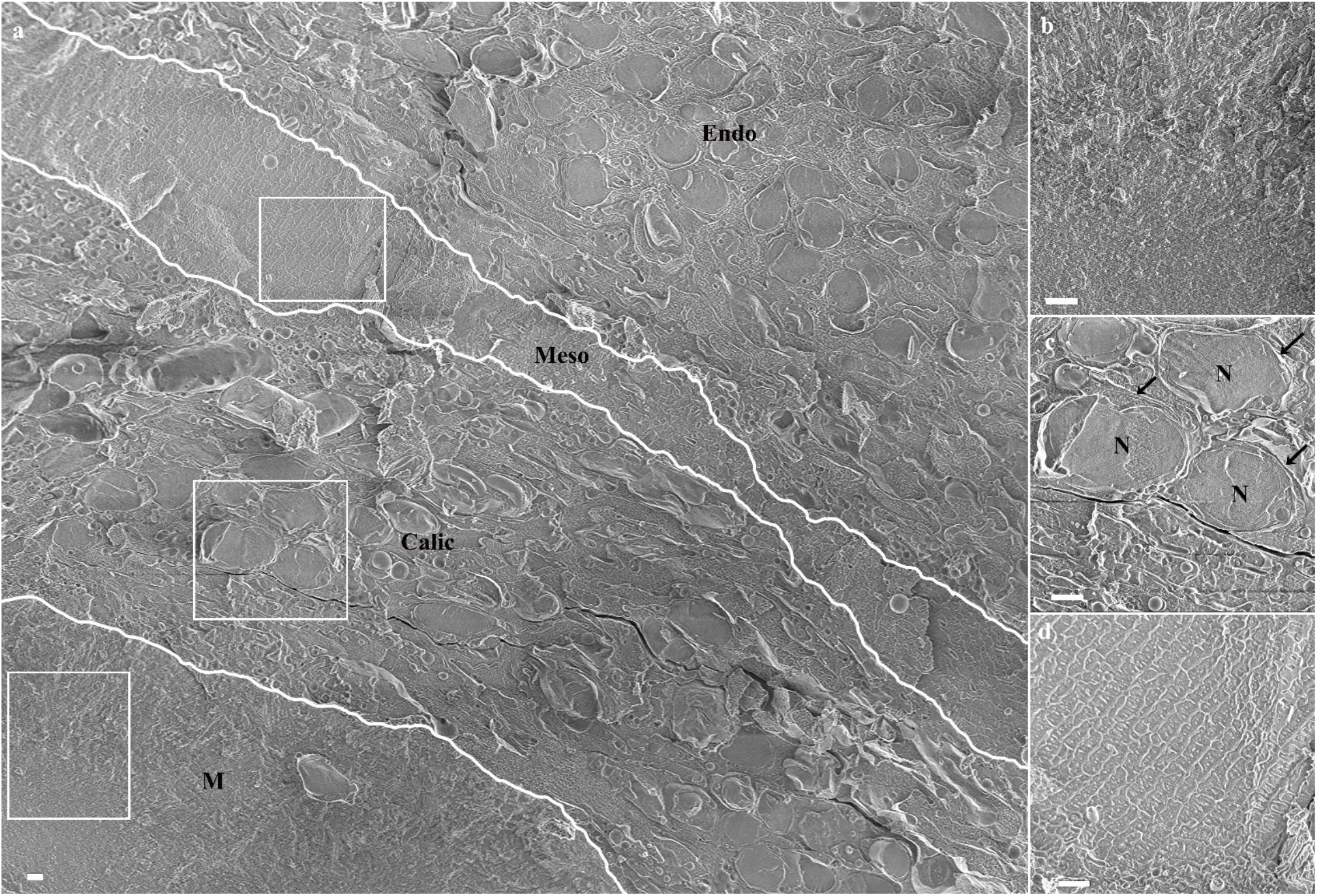
A cryo-SEM micrograph of aboral body layers observed in a high-pressure frozen, freeze-fractured primary polyp. (a) An overview image showing aboral body layers depicted by white separating lines including the mineral, **M**, (white rectangle is magnified in (b)), the calicoblastic cell layer, **Calic**, (white rectangle is magnified in (c)), the non-cellular mesoglea, **Meso**, (white rectangle is magnified in (d)) and the endoderm (**Endo**). **N**-Nucleus. All scale bars are 1µm.

The freeze-fractured surface of a septum (Fig. 3) reveals the micro-structure of the first formed mineralization zone, the CoC (Fig. 3c, f pseudo-orange), characterized by a nano-granulated surface texture composed of tightly packed nano-spheres of a uniform size of 20 ±3.1 nm (Fig. 3h). Such surface texture may implies its formation via an amorphous calcium carbonate (ACC) precursor (*44*). The remainder of the septum is composed of elongated fibrous micro-crystals arranged in a three-dimensional fan around the CoC (Fig 3c, f pseudo-yellow). The interface between the CoC and the micro-crystals layer is better observed in a newly formed septum where the CoC is already completed but the fibrous micro-crystals are just starting to form (Fig. 3 d, e, f). The fibrous micro-crystals have a flat surface and resemble single aragonite crystals as also reported in (*23*), sized up to 1 µm in width and a few microns in length, (Fig. 3f, g). The two mineral layers appear to be tightly inter-grown.

**Fig. 3:**
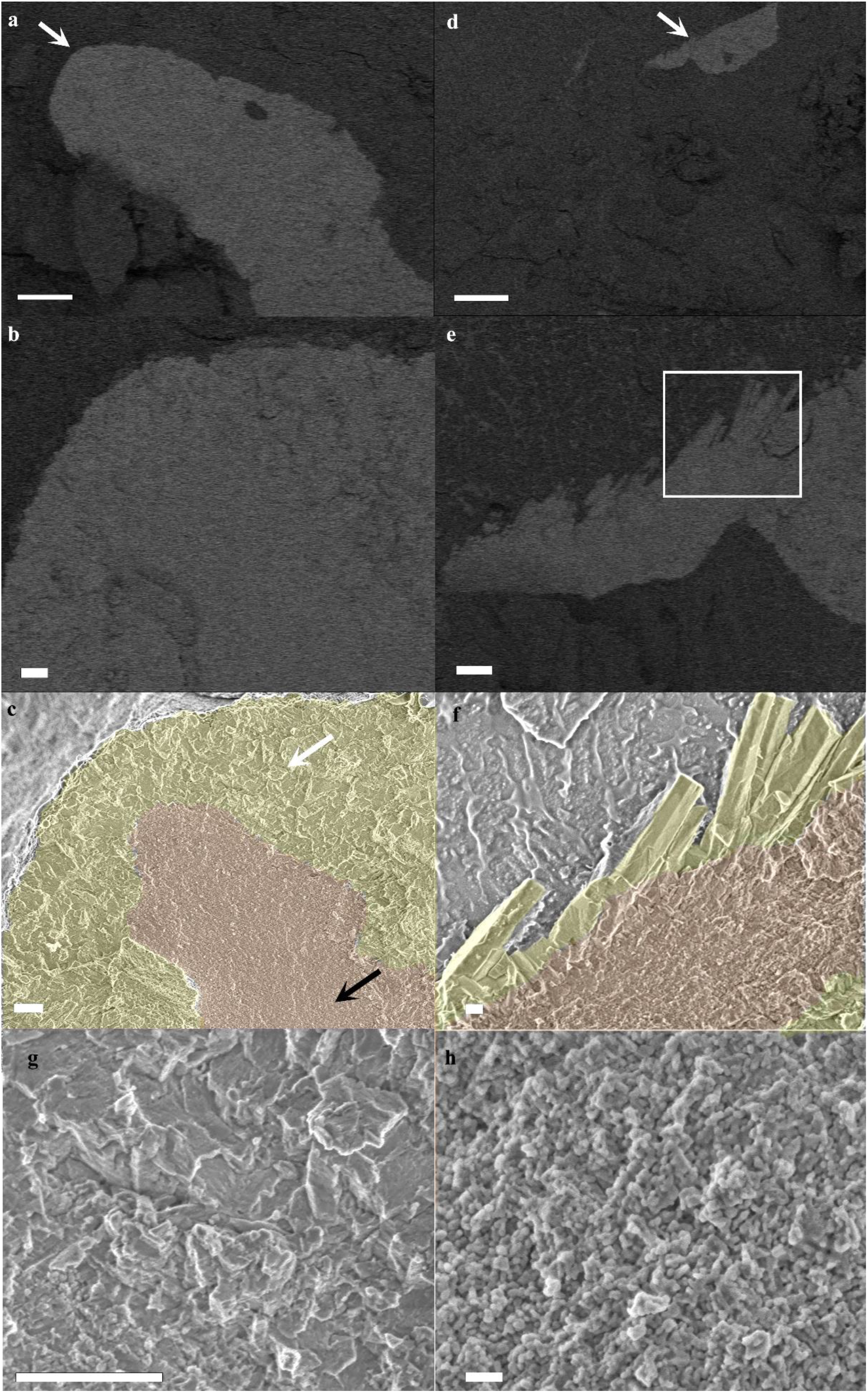
cryo-SEM micrographs of mature and forming septa in a primary polyp. (a) a mature septum imaged in ESB mode. Mineral appear brighter than its surrounding tissue. (b) Magnification of the area pointed with white arrow in (a), (c) The same areas as in (b) imaged in SE mode with CoC and elongated microcrystals highlighted by false coloring. (d) A newly formed septum imaged in ESB mode. (e) Magnification of the area pointed with white arrow in (d). (f) SE mode image of the area marked with white rectangle in (e) with CoC and elongated microcrystals highlighted by false coloring. (g) Higher magnification of micron sized crystallites fracture surface pointed with white arrow in (c). (h) Higher magnification of CoC nano-spheres texture pointed with black arrow in (c). Orange-CoC. Yellow- Elongated micro-crystals. Scale bars are (a)-and (d)- 10 µm, (b), (c), (e) and (g)-1 µm, (f)-200 nm, (h)-100 nm.

The calicoblastic cell layer shows various thicknesses, ranging between a monolayer of calicoblastic cells and a layer with a thickness of 4-5 stacked calicoblastic cells. Cell morphology in this layer varies as well (Fig. 4). Some cells of the calicoblastic layer exhibit an elliptical shape (Fig. 4a). Other cells found in close proximity with the mineral surface typically have an elongated morphology with increasing surface area on the side in contact with the mineral (Fig. 4b). Near mineral corners or sharp edges, calicoblastic cells typically have a cup shape (Fig. 4c), as also reported in earlier studies (*26*). Additionally, many of the observed calicoblastic cells have cellular extensions which appear to be a filopodia network (*45, 46*) that spans up to several cell diameters and typically occupy the space between the calicoblastic cell layer and the mineral (Fig. 4a). These filopodia are enriched with vesicles engulfed by the cell membrane (Fig. 4a,d) similar to vesicles documented in the calicoblastic layer of stony corals in previous studies, sometimes referred as secretory vesicles (*18, 47*), spherical extracellular material (*21*) or intracellular vesicles (*28*). High magnification cryo-SEM analysis of a cryo-planed primary polyp reveals the filopodia cross section and shows that the vast majority of the vesicles are found between two filopodia membranes and thus are intracellular rather than occupying the extracellular space (Fig 4d). Cryo-ESB images of the same area show that the vesicles do not have an increased electron density and therefore presumably do not contain a solid mineral or dense cation storage (Fig. 4e). Cryo-SEM/EDS analysis of calicoblastic cells found in close proximity with the septum (Fig. 5) shows that some, but not all, intracellular vesicles are enriched in carbon content compared with the cell cytoplasm and surrounding tissue. This includes both vesicles found within the calicoblastic cell body (Fig. 5b, c, d) and vesicles within filopodia (Fig. 5e, f, g). These observations were consistent across all imaged septa of six freeze-fractured or cryo-planed primary polyps.

**Fig. 4:**
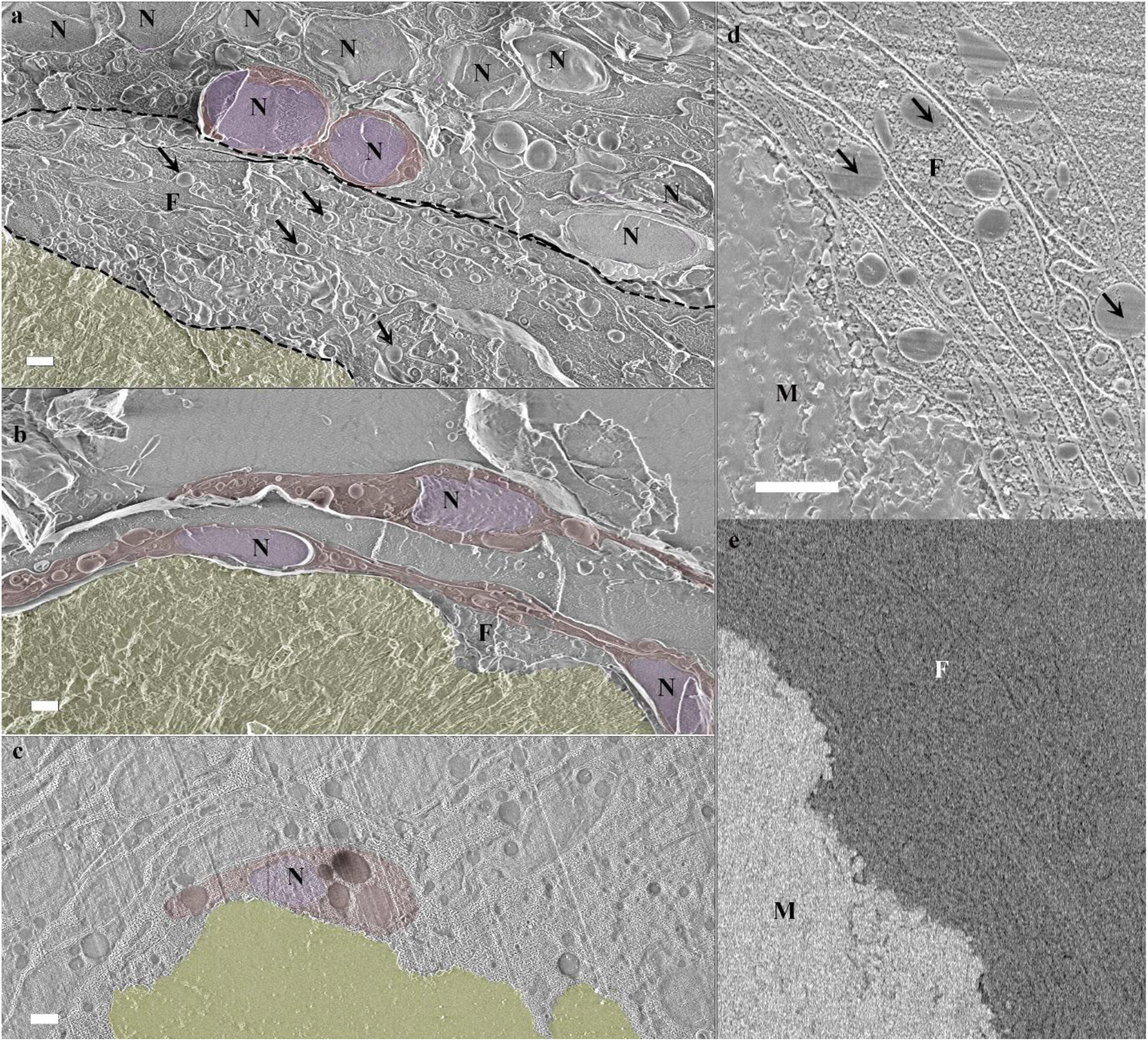
Calicoblastic cell morphologies observed in freeze-fractured (a-b) and cryo-planed (c-e) primary polyp using cryo-SEM. (a) Calicoblastic cell layer (Imaged loci is the same as in Fig. 2c) with the septum mineral and two representative calicoblastic cells highlighted by false coloring. A filopodia network found between the calicoblastic cell bodies and the septum is denoted with dashed black lines. Four representative vesicles contained within the filopodia network are denoted with black arrows. (b) Three elongated calicoblast cells found in close proximity with the mineral. (c) A cup shaped calicoblastic cell attached to the mineral surface. (d) High magnification of filopodia found in close proximity with the mineral. (e) The same field of view as in (d) imaged in ESB mode. False coloring is used to highlight: Cell nucleolus (pseudo-purple), cell body (pseudo-red), septum mineral (pseudo-yellow). Scale bars are 1 µm. **N**- nucleus, **F**- filopodia, **M**- mineral.

**Fig. 5:**
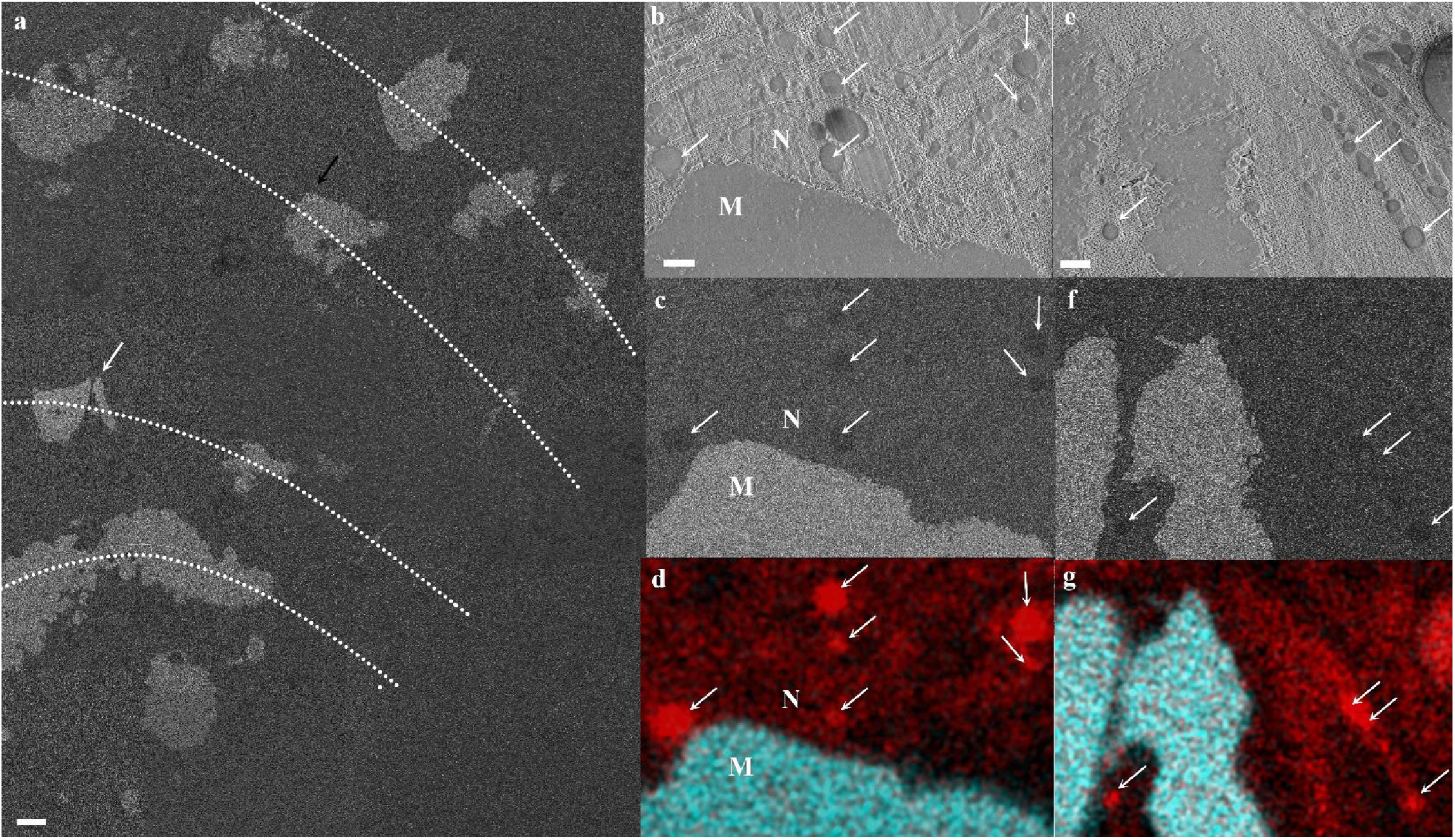
Cryo-SEM/ EDS analysis of calicoblastic cells in a cryo-planed primary polyp. (a) An overview collage image composed of several high magnification cryo-SEM (ESB mode) micrographs stitched together, showing the cryo-planed surface along several septa (Dotted line resemble septa long axis) in a primary polyp. The mineral surfaces of the planed septa appear white and the coral soft tissues appear grey. (b-d) High magnification of locus pointed with black arrow in (a) (same area as in Fig. 4c) showing a cup shaped calicoblast attached to the mineral imaged in SE, ESB and EDS map respectively. (e-g) High magnification of locus pointed with white arrow in (a) showing filopodia in close proximity with the mineral imaged in SE, ESB, and EDS maps respectively. Carbon rich vesicles are pointed with white arrows in (b)-(g) EDS maps show carbon (red) and calcium (turquoise) distributions. **M**- mineral, **N**- nucleus. Scale bars: (a)- 10 µm, (b-g)- 1µm.

### Paracellular space in the calicoblastic layer

Moving up scale from cellular morphology to tissue arrangement of the calicoblastic cell layer, we obtained a large high-resolution overview image of the micro-environment around one septum (Fig. 6a) also imaged in figure 5 (Fig. 5a-white arrow and Fig. 5e, f, and g). This overview image reveals a highly dispersed packing of calicoblastic cells (Fig. 6a pseudo-burgundy) adjacent to the septum with micrometer-sized spaces between adjacent cell bodies (Fig. 6a black dashed line). The ECM fluid occupying the space between the septum and its neighboring cells is inter-connected with these paracellular spaces (Fig. 6a pseudo-blue). Calicoblastic cells become more tightly packed moving away from the septum and into the tissue, where we observed paracellular spaces of tens of nm (Fig. 6a white arrowhead top right corner). The ECM and paracellular spaces also contain a massive filopodia network (Fig. 6a pseudo-pink) extruding tens of microns away from the calicoblastic cells bodies (Fig. 6a pseudo-burgundy) from which they are derived, towards the septum and containing a large amount of vesicles (asterisk) with an average size of 400 ±100 nm (N=100). The elaborated calicoblastic cell filopodial network we observed using cryo-SEM imaging is also correlated with the recently published scRNA-seq data of S. *pistillata* primary polyps of the same age (*37*). Analysis of the RNA-seq data shows enrichment of membrane and actin-based cell projections and transport vesicle membrane Gene Ontology (GO) terms associated with calicoblastic cells of the primary polyp. Moreover, calicoblastic cells of the primary polyps show high expression (compared with other cell types) of genes involved in filopodia network formation and function such as actins, actin bundling proteins (e.g., fascin), actin binding proteins (e.g., formins) and Arp2/3 proteins known to play essential roles in the regulation of filopodia generation (*48*) (Fig. 6b). In addition, we found relatively high expression of genes related to vesicular transport, exocytosis and the SNARE complex such as clathrin, synapto tagmins, and Ras-related proteins (RAPs)(*49, 50*) which correlates with the large amount of vesicles observed within the filopodia network (Fig. 6b).

**Fig 6:**
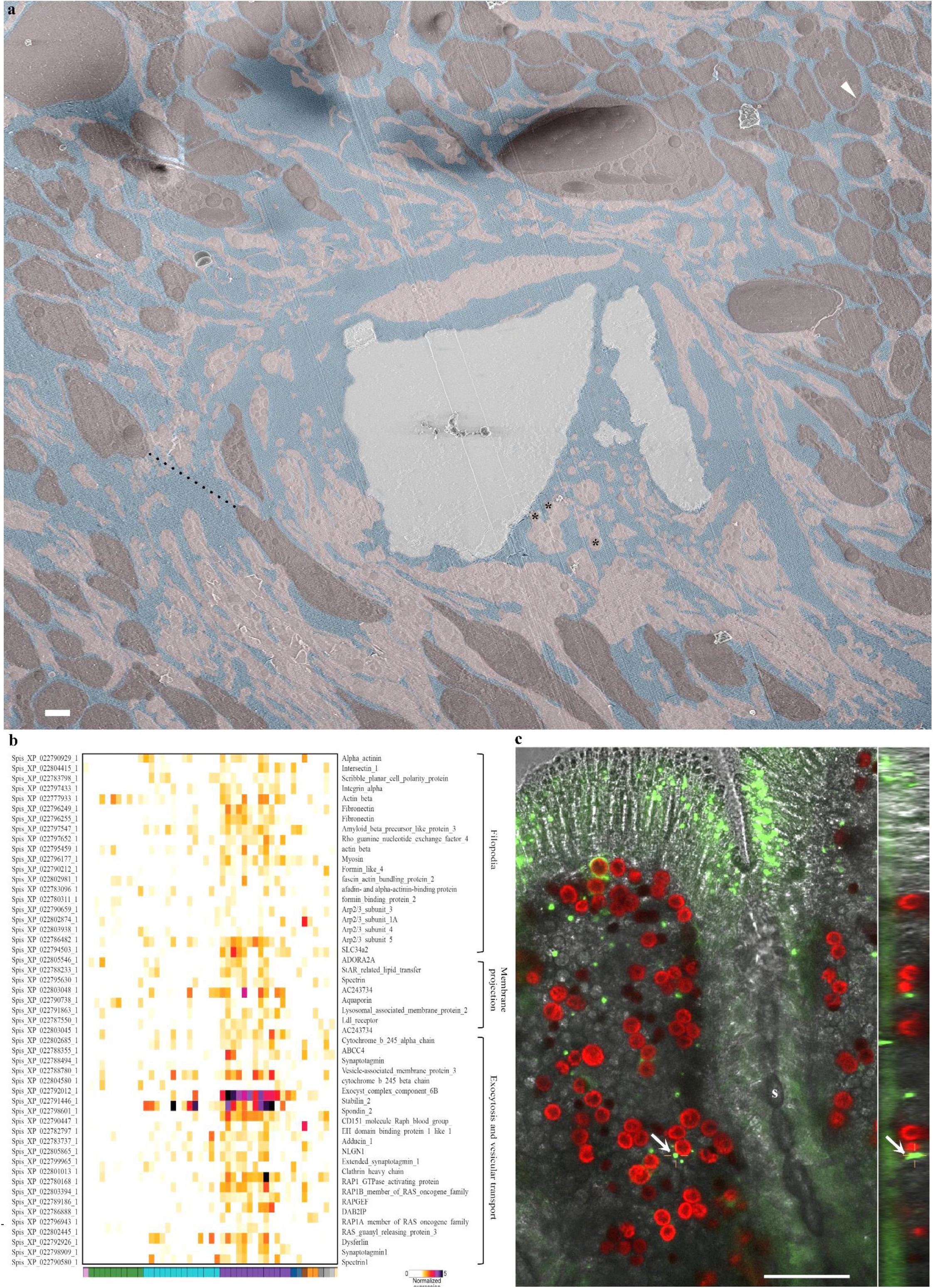
The paracellular pathway in primary polyps. (a) An overview image stitched from several high magnification cryo-SEM micrographs of the calicoblastic tissue around a septum in a cryo-planed primary polyp (same area as pointed with white arrow in Fig 5a). An area of tightly packed calicoblastic cells with paracellular space of 30 nm is pointed with white arrow head (top right corner). An area of dispersed cell packing with paracellular spacing of 8.4 µm is denoted with black dashed line. Calicoblastic cell bodies are highlighted with pseudo-burgundy, filopodia in pseudo-pink, ECM in pseudo blue and the mineral in pseudo-grey. Three representative vesicles contained within the filopodia network are marked with black astrisks. Scale bar is 2 µm. (b) Gene expression heatmap for selected genes involved in filopodia structure and function, membrane projections, exocytosis and vesicular transport, across all cell types of *S. pistillata* primary polyp based on the primary polyp scRNAseq published by Levy et al. *(37)*. (c) In-vivo confocal laser scanning fluorescence image of the tissue around a septum (s) in a primary polyp labeled with green fluorescent beads of 1µm size. Fluorescence image is composed of three channels: Green- fluorescence beads, Red- symbionts auto-fluorescence and Grey scale-transmitted laser scanning image. Center large panel is one horizontal (xy) plane taken 16µm above the glass bottom and found roughly in the middle of the z-stack data set that covers the entire thickness (30µm) of the primary polyp tissue. Fluorescent beads in this image are incorporated inside the coral tissue. The right and the bottom panels show the two side views of the z-stack data set (i.e. xz, and yz). One representative green fluorescent bead is denoted with white arrow in all three panels.

### Tissue permeability of primary polyps

Cryo-SEM imaging of cryo-planed specimens has an inherent tradeoff between high resolution and a larger field of view, which does not allow us to obtain a continuous image that follows the dimensions of the paracellular spaces or the paracellular pathway all the way from the septum to the external seawater (a distance of few hundreds µm). However, we studied the overall tissue permeability, i.e. from the external seawater inwards to the primary polyp body using fluorescent beads labeling followed by in-vivo imaging. Previous studies on adult S. *pistillata* micro-colonies show that beads of sizes larger than 20 nm do not pass through the oral epithelial layers of the micro-colonies (*42*). We therefore conducted an in-vivo labeling experiment on primary polyps using green fluorescent beads of a larger diameter (1 µm) in order to check their tissue permeability and to test our assumption that primary polyps have higher tissue permeability then adult micro-colonies based on the dispersed tissue arrangement we observed in their calicoblastic cell layer. Indeed, after only 2 hr of incubation with seawater solution containing the green fluorescent beads, the beads were observed inside the primary polyp tissue, as detected in a z-stack of the entire primary polyp obtained using in-vivo laser scanning confocal imaging (Fig. 6c). This supports the notion that tissue permeability is indeed significantly higher in the S. *pistillata* primary polyp then previously documented in corals using adult colonies of the same species (*42*).

### ECM thickness and elemental composition

We further observe that the thickness of the ECM layer is highly variable and changes in space and time in the primary polyp. In-vivo laser scanning time-lapse imaging shows a contraction and expansion movement of the ECM layer near all forming septa of the primary polyp that changes the thickness of the ECM layer facing the septum every few min (see supplementary video S3). The measured ECM thickness at one locus near the septum changes from 27 µm to 43 µm within 3 min and back to 18 µm within the next 3 min (Fig. 7a-d). This contraction movement is similar to the ECM pocket contraction movement documented in A. *digitifera* primary polyps (*51*). High-resolution cryo-SEM collage shows variable ECM layer thicknesses along a septum surface ranging from several nm and up to tens of μm (Fig. 7e). We refer to areas with a thick ECM layer bounded by loci of ECM narrowing on either side as ‘ECM pockets’ because, together with the septum surface, these areas create semi-delimited ECM spaces. We further used quantitative cryo-SEM/EDS analysis of a cryo-planed primary polyp specimen to study the elemental composition of the ECM (Fig. 8). While natural distribution of major seawater ions is well-documented in the coral skeleton using electron probe, ICP-OES, ICP-MS (*36, 52*–*55*) or dry SEM/EDS (*56, 57*), cryo-SEM/EDS is one of the few cutting edge techniques that allows in-situ detection and imaging of the distribution of these ions in the soft tissue, ECM, and skeleton at the same time. The above technique allows identification of ions in vitrified solutions down to concentrations of a few tens of mM and quantification of ion ratios in the solution (*33*). The major element in both the ECM and the cytoplasm is oxygen which is abundant in water-based solutions (*33*) (Fig. 8b, d). The cryo-EDS spectrum of the ECM also show increased levels of Na and Cl compared with the cytoplasm of calicoblastic cells in which Na and Cl are not detected (Fig. 8b, e and f). The latter is expected due to the fact that cellular concentrations of Na (12 mM) and Cl (10 mM) (*58*) are below the detection limits of this technique which are 25-50 mM and 73-50 mM, respectively (*33*). The mineral is easily differentiated form the soft tissue by its strong Ca signal (Fig. 7c). Increased levels of Na are also observed in the mineralized septum (Fig. 8f) unlike Cl levels which appear very low in the mineral (Fig. 8e). This is because Na is incorporated into the aragonite lattice of coral skeleton with a molar ratio of 1-2% (Na/Ca) (*53, 54*). We used the cryo-EDS analysis to obtain the Cl:Na ratio in the ECM which is 1.08±0.05, and is thus close to the Cl:Na ratio in seawater, i.e., 1.13 (*59*), and different from the cellular Cl:Na ratio, 0.8 or lower (*58*). After Na and Cl, the next major seawater ions are Mg^2+^, SO_4_^2-^, Ca^2+^ and K^+^. These cations have natural concentrations in seawater that are below the detection limit of the cryo-EDS technique (*33*), and therefore are not detected in the ECM in our analysis. This is, to our knowledge, the first direct imaging of dissolved Na and Cl in the ECM solution. These results strongly support that the ECM contains seawater.

**Fig. 7:**
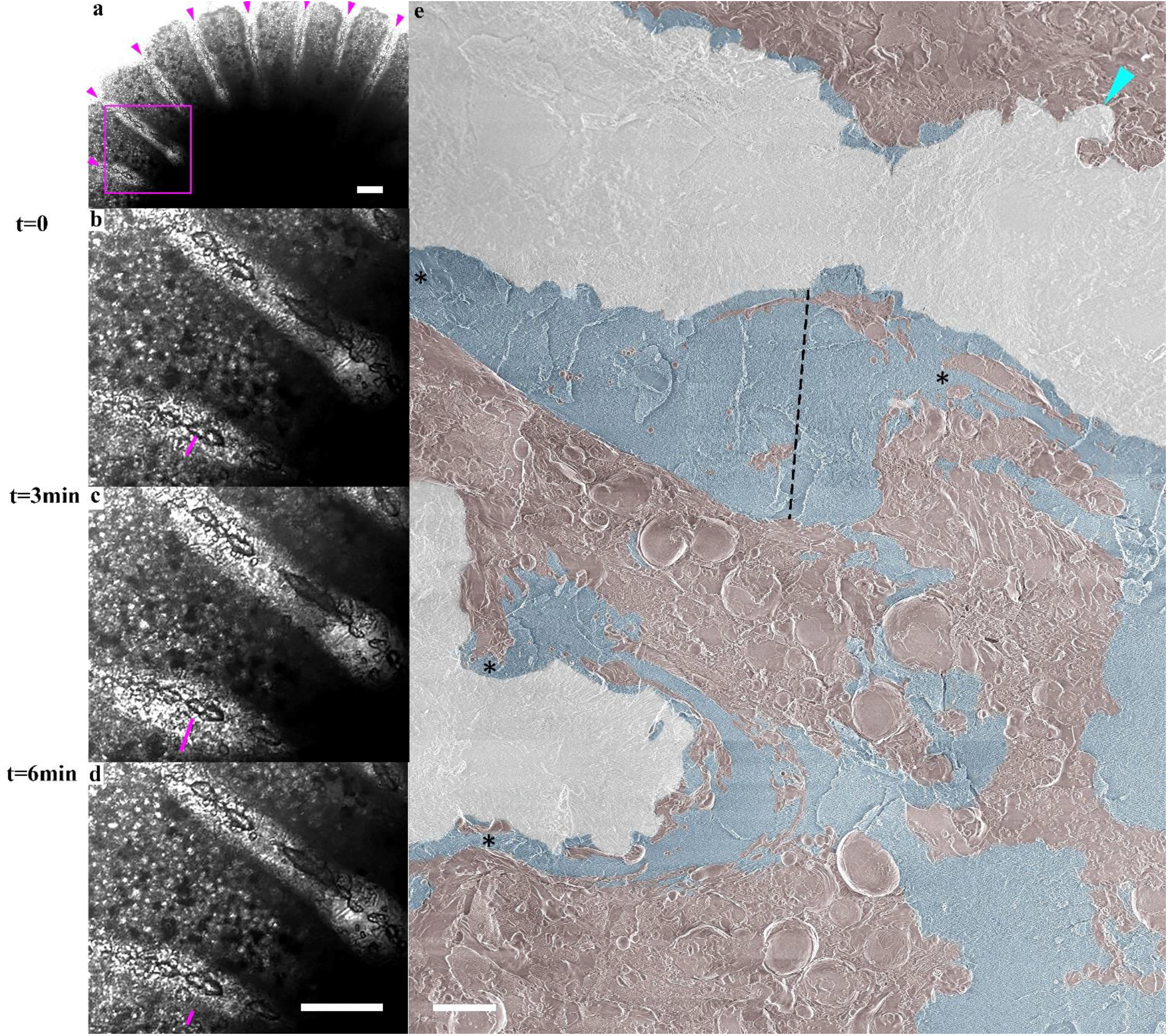
ECM thickness variation along primary polyp septa. a) Transmitted laser scanning in-vivo microscopy image of a primary polyp with forming septa denoted with magenta arrowheads recorded at t=0. (b)-(d) Higher magnification of the area marked with magenta square in (a) recorded at t=0, t=3 min and t=6 min respectively (see also supplementary video S3). The thickness of the ECM on the bottom side of one septum is marked with magenta line and equals 27 µm, 43 µm and 18 µm respectively. (e) A collage of high resolution cryo-SEM micrographs stitched together to form and an overview image of the a freeze-fractured septa of a primary polyp. Mineral surface is highlighted in pseudo-grey, the ECM in pseudo-blue, and the coral tissue including cell bodies and filopodia are highlighted in pseudo-burgundy. The thickness of the ECM layer in one ‘ECM pocket’ is depicted by dotted black line and equals-38 µm. Areas of ECM layer narrowing are pointed with asterisks. An area where tissue is closely attached to the mineral surface is pointed with turquoise arrowhead. Scale bars: (a-d): 100 µm, (e): 10 µm.

**Fig 8:**
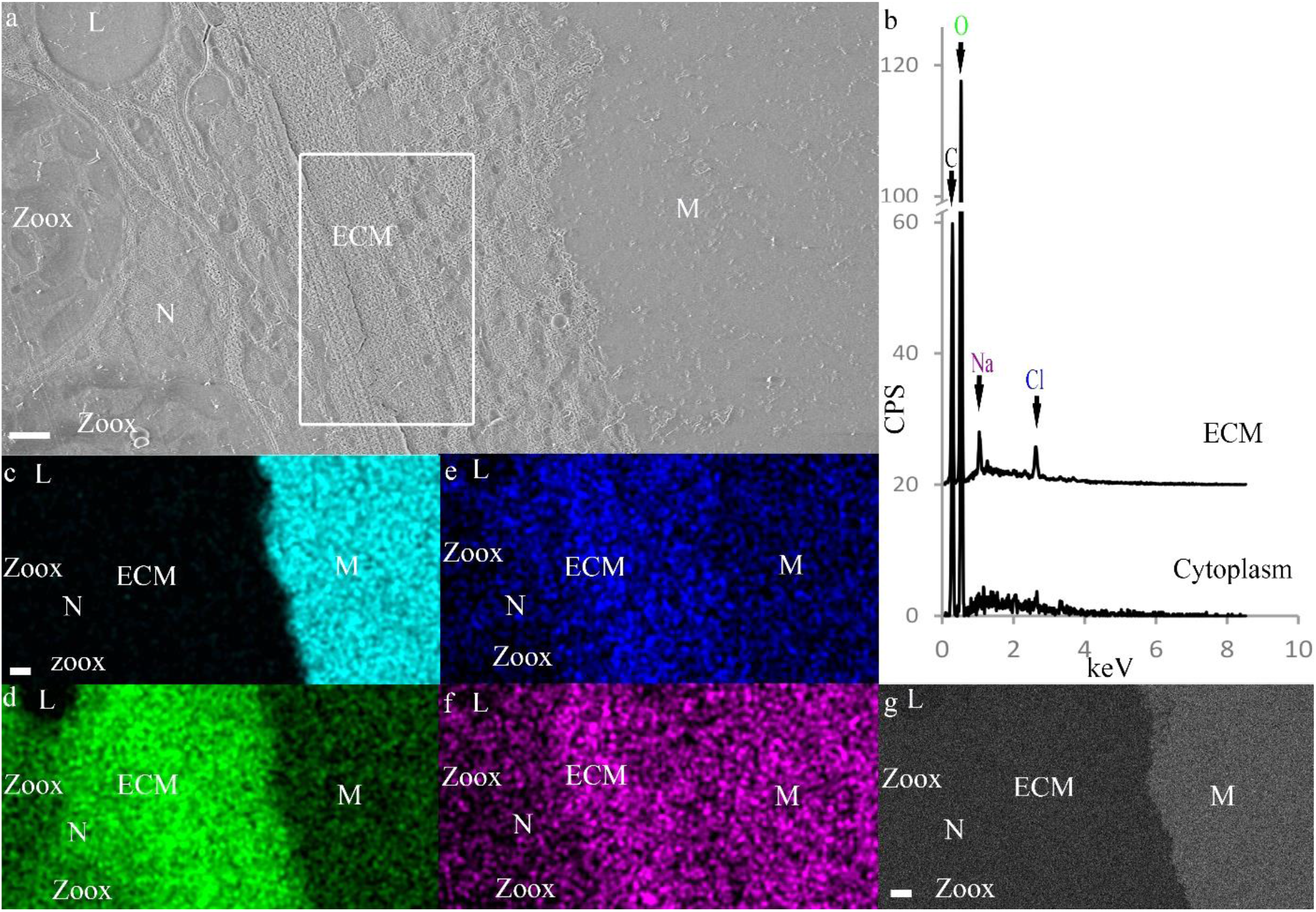
Cryo-SEM/EDS analysis of the ECM. (a) SE mode micrograph of the tissue-mineral interface. White rectangle marks ECM area enclosed between the mineral and the calicoblastic layer. (b) Cryo-EDS spectra of the ECM (measured area is the area marked with white rectangle in (a)) and of the cytoplasm in a calicoblastic cell. Identified elements are labeled in the graph. (c)-(f) Cryo-EDS elemental distribution maps of the same area of image (a) of the elements: Calcium (Cyan), oxygen (Green), chlorine (Blue) and sodium (Magenta) respectively. (g) Cryo-ESB micrograph of the same area as in (a)-(f). All scale bars are 1 µm. M= Mineral, N= Nucleolus, Zoox= algal symbiont, L= Lipid body. CSP= counts per second.

## Discussion

We combined in vivo dynamics, morphological characterization and elemental analysis together with scRNA-seq analysis, techniques coming from two different fields of study, to characterize the calcifying interface during skeleton formation in the primary polyp life stage of the stony coral *S. pistillata*. Our cryo-SEM observations confirm the mineral micro-structure, the aboral tissue layers and calicoblastic cell morphologies previously documented in stony corals using other imaging techniques (*4, 22, 24, 26*). We further show that calicoblastic cells produce an elaborate filopodial network containing a multitudinous population of carbon-enriched vesicles that processes towards the tissue mineral interface. The large dimensions of this network with respect to the size of the calicoblastic cells, as well as the related up-regulation in calicoblasts of genes associated with filopodia development, imply that these networks play an important role in calicoblastic cell activity. We also documented unique tissue arrangements at the mineral-calicoblastic layer interface which include larger ECM pockets and paracellular spaces than previous estimations (*4, 26, 28*). This goes together with our observation of high tissue permeability in the primary polyp, and, with our indication of seawater in the ECM, as suggested in previous studies (*51, 52, 60*). One possible function of the increased tissue permeability and large ECM volumes composing seawater documented here in primary polyps may be to enhance ion transport for mineralization in order to support high mineralization rates in this life stage, as also supported by the scRNA-seq data (*37*).

### Seawater transport to the mineralization site

Indications for a paracellular pathway connecting the external seawater with the mineralization site were found in stony corals by using the cell impermeable dye calcein (*42, 51, 61, 62*). Further indication for the transport of seawater to the mineralization site comes from stable isotope incorporation experiments (*52*). In this study, we used cryo-EDS analysis to directly image the ECM fluid and its elemental composition in a ‘native like’ state and found further evidence that seawater is indeed incorporated into the ECM. This brings up intriguing questions such as: (i) What is the volume and turn-over rate of seawater in the mineralization site? and (ii) what is the role of seawater in the mineralization process? (*4, 22, 42, 52*) Two important parameters required to pursue these questions are the permeability of the coral tissue and the dimensions of the ECM layer. Tissue permeability tests performed on adult *S. pistillata* micro-colonies showed that molecules and particles of sizes between 13 Ȧ-20 nm diffuse via the paracellular pathway; this size range was therefore considered as the size of the intercellular junctions connecting one cell to the other, i.e. the coral ‘septate junctions’ (*42*). Indeed, septate junctions have been documented and characterized in *S. pistillata* micro-colonies (*26, 63*). However, observations of the present study show a significantly higher tissue permeability of primary polyps compared with their adult counterpart, with particles of 1 µm size passing through their epithelial tissue (Fig. 6c). Two possible pathways for particle incorporation into tissues are an intracellular pathway (micropinocytosis) and a paracellular pathway. A size of 1 µm is much larger than the maximal size of particles documented to be incorporated into the coral epithelial tissue via macropinocytosis, i.e., 200nm (*62*), although this does not ruled out micropinocytosis as an incorporation pathway at this stge. A size of 1 µm also cannot be attributed to a septate junction (*63*). Nevertheless, in order for septate junction size to impose whole tissue permeability, an assumption must be made, that all adjacent cells are attached to one another via septate junctions. The dispersed cell packing in the calicoblastic cell layer with paracellular spaces of few micrometers (Fig. 6a) documented here, challenges this basic assumption. We therefore infer that while some calicoblastic cells are attached to their neighboring cells via septate junctions, others are bathed within the ECM and separated by few microns from some of their adjacent cells. Our observations, thus, show that tissue permeability is not solely defined by septate junction dimensions, and that it is significantly higher in primary polyps than previously documented for adult micro-colonies of the same species (*42*). We describe here an intriguing and highly dispersed cellular arrangement of the calicoblastic layer, contradicting the previous conceptual framework on cell packing in the coral calicoblastic tissue and possibly even in epithelial tissues of other organisms. One reason why the dispersed cell packing documented here was not reported in earlier studies performed on *S. pistillata* may be that it is more characteristic of the primary polyps then of the adult life stage which was mainly used in previous studies (*26, 28, 47*). Additionally, the loosely attached cells found in close proximity with the septa even of existing in adult specimen are expected to be underrepresented in TEM thin sections, histological sections and fluorescent imaging involving a post-fixation demineralization procedure (which are the major techniques used in previous studies). This is because the loosely attached parts of the tissue can be washed away from the sample during de-mineralization and solution exchange steps involved in sample preparation (*28*). Therefore, the possible existence of a dispersed cell packing also in adult corals should not be rule out.

We also observe that cellular packing can vary between dispersed and tight arrangements at different locations within the same coral specimen. It is as yet unclear whether the observed differential arrangement is actively controlled. One function of a dispersed cellular arrangement may be used to locally and temporally change tissue permeability. Corals were shown to control their overall body permeability and to modify it in response to external stressors such as osmotic pressure and temperature change (*61*). It is, therefore, reasonable to assume that they can also locally modify their tissue permeability along the forming skeleton according to their needs. One possible role of the paracellular pathway connecting the external seawater with the ECM is the transport of ions used for mineralization to the mineralization site. In such a case, changing tissue permeability may be a way to increase seawater supply to the mineralization site during periods of rapid mineralization activity. This is also while increasing cell surface area in contact with seawater, which may be used to increase absorption of ions from the seawater into the cells. However, any relation between locally increased tissue permeability and the mineralization activity has yet to be demonstrated in stony corals. Regardless of the function that tissue permeability modification plays in stony corals, the observation that primary polyps have higher tissue permeability than adult coral colonies may make them more vulnerable to micro-plastic contamination, sediment suspension or sewage pollution and should, therefore, be further studied and taken into account in coral reef management.

Another important parameter to understand the role of incorporated seawater in the calcifying space, in addition to coral tissue permeability, is the thickness of the ECM layer. The ECM is the site where aragonite micro-crystals comprising the coral skeleton crystalize and grow (*4, 51*). The ECM is largely documented as a thin non-cellular fluid or gelatinous layer with a thickness of nanometers to 1 micron filling the space between the mineral and the calicoblastic tissue (*4, 22, 24, 26, 28*). Previously hypothesized (*24, 25*) areas in which calicoblastic tissue is locally lifted away from the mineral creating semi-delimited ECM spaces are referred as ‘ECM pockets’. Recent studies using in-vivo fluorescence imaging show growing evidence for such pockets and further report tissue contraction movements in these pockets that facilitate flow of the ECM fluid between them (*51*). Our in-vivo observation also support simultaneous contraction movements of the tissue around the forming septa, which modifies the ECM layer thickness along them in primary polyps. Exact measurements, however, of the thickness of the ECM layer in these pockets cannot be made solely based on light microscopy, due to resolution, optical contrast and penetration depth limitations, or either based on histological sections or TEM thin sections in which the ECM fluid is effectively replaced during preparation procedures. We thus used cryo-fixation (keeping the sample fully hydrated) and cryo-SEM imaging to measure the local thickness of the ECM layer. We documented a thickness of tens of µm of the ECM in such pockets, which is significantly larger than previous estimations (*4*). We also document high variability in ECM thickness measured along septal surfaces. These findings are relevant to achieve a better understanding of the biomineralization mechanism taking place in the ECM layer. One model discussed in the literature is ion-by-ion crystallization of the aragonite micro-crystals from a saturated solution, where the ECM fluid functions as the saturated mother solution for mineralization (*64*). Stable isotope incorporation studies (*52*) support the ion-by-ion strategy. This strategy requires the cycling of large volumes of the mother solution for mineralization in the organism body. A missing link in this model for calculating seawater turn-over rates in the ECM is the ratio between the volume of the ECM and the surface area of the septum mineral, i.e. the ECM thickness considering a simplified box shape of the ECM. The model predicts ECM thickness of tens of µm, and therefore observations of the current study supports the feasibility of this model (*52, 55*). However, our observations of a high spatial and temporal variation of the ECM thickness in the current study imposes more complexity on the calculations than using a simplified box model with a constant ECM thickness. Spatial variability of the ECM thickness along the septum suggests active tuning of the ECM size and shape according to mineralization needs. This is in agreement with previous observations that coral mineralization is non-continuous along the skeleton but is rather patchy in time and space on a spatial scale greater than tens of microns (*52, 65*).

The observation that ECM pockets can reach up to tens of microns in thickness also helps to explain another recent observation of primary cilia in some calicoblastic cells of *S. pistillata* micro-colonies (*66*). Primary cilia are differentiated from filopodia by their shorter length and straight stalk-like morphology. They also make up a much smaller portion of the surface area of cells, as they are restricted to one primary cilium per calicoblastic cell. While primary cilia, like filapodia, can be observed by cryo-SEM as used in the current study, it is hard to differentiate them from other cytoplasmic extrusion using this technology alone. Primary cilia act as mechanosensors that translate extracellular stimulations from the external micro-environment to intracellular signals in different organisms (*67*), and in the case of corals they are thought to transfer signals from the ECM to the calicoblastic cells (*66*). One question raised by those authors is whether the cilia have enough space in the ECM to stretch and bend considering their length of 1-2 µm. Our observation of the range of ECM thicknesses shows that they certainly do. It is possible, therefore, that primary cilia play a role in sensing and controlling ECM fluid flow inside ECM pockets.

The high-pressure freezing fixation technique used in this study holds limitation on specimen dimensions, which are up to 3mm in diameter and 200µm in thickness (*29*). Thus, allowing the analysis of primary polyps but not of adult corals. The observation of increased tissue permeability and thick ECM pockets composing seawater may, therefore, be attributed only to primary polyps at this stage. However, possible manifestation of these tissue arrangements also in the adult life stage should not be ruled out. The calicoblastic filopodia network composing large population of carbon rich vesicles observed at the interface between the cells and the forming skeleton is intriguing and require further study in order to resolve its function. We also cannot deduce the contents of the vesicles with the techniques used in this study. Carbon enrichment in these vesicles may be attributed to skeletal organic biomolecules transported to the mineralization site to construct the forming skeleton, but may also be attributed to other organic molecules used for different physiological processes carried out by the calicoblastic cells other than the mineralization process.

### Ions for mineralization

While we provide here further evidence that seawater is a component of the ECM, it is important to stress that this does not mean the ECM directly reflects ocean chemistry or that external changes in seawater chemistry affects skeleton mineralization in an uncontrolled manner. Previous studies and our cryo-SEM observations reported here show that ECM pockets are largely delimited spaces, in which internalized seawater is prone to strict biological control and modification induced by the calicoblastic cells. The control and alteration of ECM chemistry by the coral tissue is clearly evident from previous work showing elevated concentrations of Ca^2+^, CO_3_^2-^, pH levels and thus aragonite saturation state (Ω_arag_) inside ECM pockets relative to the external seawater (*68, 69*). The concentration of Ca^2+^ ions in seawater is roughly fifty times higher than CO_3_^2-^ concentration (*69*). Therefore, upon delivery of external seawater to the mineralization site (as supported by our observations) CO_3_^2-^ is the limiting factor for mineralization, although there is some evidence that HCO_3_^-^ also contribute to the DIC pool used for mineralization (*70*). This is in agreement with stable isotope incorporation studies suggesting that the major part of Ca^2+^ ions used for coral biomineralization is delivered as dissolved Ca^2+^ ions found in the seawater which is incorporated into the ECM, rather than active Ca^2+^ pumping (*52*). Moreover, It has been consistently observed by-proxy and by direct micro-electrode measurements that both CO_3_^2-^ and total dissolved inorganic carbon (DIC) levels are elevated above seawater levels in the ECM (*52, 69*–*73*). The scRNA-seq data of the primary polyps may also support a possible DIC concentration mechanism. The latter shows that two bicarbonate co-transporters (SLC4γ and SLC4A10) are among the most specific and highly expressed genes in the calicoblasts of primary polyps, with more than 80% of their total expression in calicoblast. SLC4γ that was reported to be a specific isoform to stony corals (*74*), shows even more specific expression, with 90% of its total expression in the calicoblastic cells (Fig. S1). These results are consistent with previous studies showing a specific immunolocalization of SLC4γ to the calicoblastic ectoderm, with the authors suggesting SLC4γ to be responsible for supplying bicarbonate to the calcification site (*74*). In addition, according to the scRNA-seq data, primary polyp calicoblasts show enrichment of six carbonic anhydrase (CA) genes, encoding enzymes that catalyze the interconversion of carbon dioxide and bicarbonate (Fig. S4), including STPCA (XP_022801446) shown to be localized to calicoblasts (*75*) and STPCA2 (XP_022799914) found in *S. pistillata* skeleton proteomes (*39, 76*).

While some evidence support an ion-by-ion crystallization strategy exploited in stony corals skeletogenesis (*52*), other observations support an alternative strategy of crystallization via an amorphous calcium carbonate (ACC) precursor phase (*57, 71, 77, 78*). In this study, we did not observe any dense Ca^2+^ storage compartments in the calicoblastic cell layer, the ECM or in any of the other coral tissue layers. Ca^2+^ concentrations in an ACC phase are typically in the molar range (*79*) and therefore well within the detection limit of the cryo-EDS technique, which is 25-50 mM (*33*). However, this negative observation does not rule out the existence of such phases in other parts of the primary polyp body, in sizes smaller than our resolution limits (few nm) or in times points or life stages other than that of the primary polyps studied here. Mineralization via an ACC precursor may also be more pronounced in the formation of the CoCs, which are characterized by a nano-granulated surface texture typical of biominerals formed via an ACC precursor (*44*), compared with the elongated micro-crystallites composing the rest of the septum (*23*). Indeed, a bi-model combining both mineralization via an ACC precursor and ion-by-ion mineralization from a saturated solution has been proposed in the literature (*71*).

We used a combined structural, chemical and molecular analysis approach to gain new insights into the biomineralization process of stony corals, a process integral to the health, resilience, and persistence of coral reefs. Our observations clarify the range of dimensions between cells, and of the calcifying space, i.e. between the tissue and the skeleton and reveal increased tissue permeability in the primary polyp life stage compared with the tissue permeability of the adult life stage documented in previous works. These measures are important links that were previously missing in efforts to rectify the different biomineralization models debated in the literature. We show an intimate involvement of large volumes of incorporated seawater in the mineralization site, thus providing a new conceptual framework for understanding the ‘vital effect’ observed in geochemical studied using paleoceanographic tracers contained in stony coral skeletons (*16, 52, 60*). These observations are also important for better understanding the resilience of newly recruited primary polyps to different stressors affecting the coral reef. The increased tissue permeability and rapid incorporation of the external seawater to the mineralization site potentially make these newly recruited corals more vulnerable than adult corals to changes in the external seawater. This includes current and future proposed global changes such as ocean acidification, for which corals need to compensate in order to maintain a micro-environment favoring mineralization, and local stressors such as micro-plastic contamination, sewage pollution and sediment suspension, all of which would potentially be more rapidly incorporated into the body of primary polyps compared with their adult counterparts. This should be considered in risk assessment and management of coral-reef ecosystems around the globe. By exploiting new experimental approaches, our observations add up to several recent studies (*15, 37, 68*), revealing a larger and more versatile biological toolkit exploited by corals for their biomineralization process than previously recognized.

## Materials and Methods

### Stylophora pistillata primary polyps

*S. pistillata* larvae were collected from colonies at depths of 8-14 m in the reef adjacent to the Interuniversity Institute of Marine Sciences (IUI), 29°30′06.0″N 34°54′58.3″E, in the Gulf of Aqaba (Israel) under a special permit from the Israeli Natural Parks Authority. Collection was performed using larvae traps made of 160 μm plankton nets following Neder et al. (*57*). Collected larvae were acclimated overnight under ambient conditions (∼25 °C and ∼pH 8.2) in a flow-through outdoor aquarium exposed to natural lighting with fresh seawater filtered to 60 µm. After undergoing metamorphosis, planula were allowed to settle as primary polyps using seawater volume limitation for a few minutes on a glass bottom dish (for light microscopy) or on a high-pressure freezing aluminum disc, (for cryo-fixation and cryo-SEM imaging). After initial attachment to the substrate, primary polyps were re-immersed in larger seawater volumes. The settled primary polyps commenced mineral deposition and were used for all experiments two to five days post-settlement.

### In-vivo calcein labeling and imaging

Primary polyps were incubated in freshly filtered (0.2 μm) seawater with 3 μM calcein blue (Sigma–Aldrich 54375-47-2) for 4-5 hours. Specimens were then rinsed with filtered (0.2 μm) seawater. Calcein labeled primary polyps were observed using an inverted confocal laser-scanning microscope (Nikon A1R) with a Plan Fluor 10x DIC L objective. Images were acquired in three channels: Blue (calcein, ex: 406 nm, em: 450 ±50nm), green (host endogenous green fluorescent protein, ex: 492 nm, em: 525 ±50 nm) and red (photosymbiont chlorophyll, ex: 492 nm, em: 700 ±75 nm). Pinhole size was 21.7 µm. White light images of primary polyps were also obtained using both a Leica DM2000 micro-system with a 10x Leica HI PLAN 10x/0.25 objective and Nikon eclipse T1 microscope with a color camera Nikon Ds-Ri2 using an S Plan Fluor ELWD 40x DIC N1 objective. All images were acquired with the Nikon Nis-Elements software (Nikon Instruments, Melville, NY, United States).

### In-vivo fluorescence bead labeling and imaging

Fluorescent bead labeling solution was prepared by diluting 2µl aqueous suspension of fluorescent yellow-green beads (Diameter=1µm) (Merck L4655) in 8ml freshly filtered (0.2 μm) seawater. Primary polyps were incubated in the fluorescence bead labeled seawater solution for 2hr, and imaged with inverted confocal laser-scanning microscope (Nikon A1R) with both Plan Fluor 10x DIC L and Plan Fluor 40x Oil HN2 objectives. Images were acquired in three channels: Green (fluorescence beads ex: 492 em: 525±50), red (photosymbiont chlorophyll, ex: 492 nm, em: 700 ±75 nm) and laser transmitted image. Pinhole size was 61.3 µm. Time-lapse datasets were obtained using Plan Fluor 10x DIC L objective by acquiring 49 time points in 6min. Z-stack data sets were acquired using Plan Fluor 40x Oil HN2 objective to cover the entire thickness of the primary polyp tissue in the observed area with 2µm steps. All images were acquired with the Nikon Nis-Elements software (Nikon Instruments, Melville, NY, United States).

### Cryo-EM techniques

Primary polyps 4-5 days post settlement underwent for high-pressure freezing, freeze fracture, cryo-planing and cryo-SEM/EDS imaging. The above techniques were performed following Mor Khalifa et al. (*32, 33*). In short:

#### High-pressure freezing (HPF)

Calcein labeled and untreated live 4-5 d old S. *pistillata* primary polyps were immersed in a filtered (0.2 μm) natural seawater solution containing 10 wt% dextran (Fluka) as a cryo-protectant, and immediately high-pressure frozen (HPM10, Bal-Tec AG, Liechtenstein or EM ICE, Leica Micro-systems, Vienna, Austria) between two aluminum discs. The mounting procedure took up to 30 sec. Samples for freeze fracture were frozen between two identical aluminum discs (diameter = 3 mm, thickness = 100 µm) and samples for cryo-planing were frozen inside an aluminum disc of diameter = 3 mm, thickness = 200 µm with a flat aluminum cover.

### Cryo-planing

High-pressure frozen samples were transferred to a cryo-microtome (UC6, Leica Micro-systems, Vienna, Austria) and planed at -150 °C in a nitrogen atmosphere to achieve a flat cross section surface using a diamond blade (Cryotrim 20, DIATOME, Biel, Switzerland). Samples were then vacuumed to 5 × 10^−7^ mbar, -120 °C (BAF 60, Leica Micro-systems, Vienna, Austria) and transferred for cryo-SEM imaging using a vacuum cryo-transfer device (VCT 100, Leica Micro-systems, Vienna, Austria). Finally, the samples were loaded into a scanning electron microscope, (Ultra 55 SEM, Zeiss, Oberkochen, Germany) where they were imaged at -120 °C.

### Freeze-fracture

High-pressure frozen samples were vacuumed to 5 × 10^−7^ mbar at -120 °C and a fracture was conducted at the interface between the two HPF discs containing the sample (BAF 60, Leica Micro-systems, Vienna, Austria). One disc containing the exposed cryo-fixed, freeze-fractured primary polyp was transferred for cryo-SEM imaging using a vacuum cryo-transfer device (VCT 100, Leica Micro-systems, Vienna, Austria) and loaded into the SEM where it was imaged at -120 °C.

### Cryo-Scanning Electron Microscopy /Energy Dispersive Spectroscopy (cryo-SEM/EDS) analysis

High-pressure frozen cryo-planed or freeze-fractured *S. pistillata* specimens were loaded into the SEM at -120 °C. Cryo-planed samples were then heated to -105 °C for 3-10 min (etching) to remove adsorbed surface nano-ice crystallites deposited on the sample surface during sample transfer. Freeze fractured samples did not undergo an etching procedure. Samples were then imaged to find loci of interest using an in-lens (in the column) secondary electron (SE) detector and an in-the-column energy selective backscattered electron (ESB) detector using the following microscope conditions: working distance = 2 mm, acceleration voltage = 1.5 kV and aperture = 10 µm. After loci of interest were found and imaged in high resolution using both detectors, cryo-planed samples were transferred using a vacuum cryo-transfer device (VCT 100, Leica Micro-systems, Vienna, Austria) to a freeze fracture device (BAF 060; Bal-Tec) for carbon coating (8 nm) before EDS analysis. Samples were transferred back to the electron microscope and cryo-SEM/EDS analysis was performed using the following microscope conditions: working distance = 7 mm, acceleration voltage = 9 kV, aperture = 30 µm. Cryo-EDS analysis was performed using a Bruker Quantax microanalysis system with an XFlash®6 60 mm detector. Element distribution maps and EDS spectra of areas of interest were obtained and analyzed using Esprit software.

### Cryo-fluorescence imaging

Vitrified fractured primary polyp specimen were transferred from the SEM under a cryogenic temperature via VCT (VCT 100; Leica Microsystems) to the freeze-fracture device (BAF 060; Bal-Tec) where the VCT was vented with cold gaseous nitrogen. Samples were then unloaded into liquid nitrogen and transferred into a cryo-Correlative Light Electron Microscopy (cryo-CLEM) stage (Linkam, model CMS196), pre-cooled to liquid nitrogen temperature. The cryo-stage was mounted on an upright Leica TCS SP8 MP microscope, equipped with an external Non Descanned Detectors (NDD) HyD and Acusto Optical Tunable Filter (Leica micro-systems CMS GmbH, Germany) and internal HyD detectors for confocal imaging. Second Harmonic Generation (SHG) signal was excited by a Tunable femtosecond laser 680-1080 Coherent vision II (Coherent GmbH USA). A z-stack of fluorescence images was obtained using a 10x Leica HI PLAN 10x/0.25 objective and collected in three channels: Blue (calcein, Em: 412-473), Green (host endogenous green fluorescent protein, Em: 501-551nm) and Red (photosymbiont chlorophyll, Em: 650-737nm). The xy dimension of the overview image was composed of four fields of view automatically stitched together to cover the entire specimen and the z-stack vertical range was chosen to cover the entire topographic range of the specimen fractured surface (150 μm) with a step size of 4 μm. Cryo-fluorescence images were produced by max-projection of all z-stack images using FIJI and Leica SP8MPImage analysis.

### Image analysis

We used Adobe Photoshop for brightness and contrast level adjustments of cryo-SEM/ EDS/Fluorescence micro-graphs and manual stitching of high-resolution cryo-SEM micro-graphs to obtain large field-of-view high-resolution collage images. We conducted false coloring of SE mode cryo-SEM micro-graphs to highlight identified loci of interest (raw images without false coloring can be found in supplementary material Figure S5). Microsoft Excel was used for graphical representation of EDS spectrum.

### Single cell RNA-seq data analyses

Gene expression analysis and heatmaps were created from the recently published interactive Shiny (*80*) application “https://sebe-lab.shinyapps.io/Stylophora_cell_atlas/” based on the S. *pistillata* single cell RNA-Seq (*37*). Gene expression levels as fold-change were normalized by computing a regularized geometric mean within each metacell and dividing this value by the median across metacells.

## Acknowledgments

We would like to thank Dr, Neta Varsano for the illustration in figure 1a-d; Dr. Yoseph Addadi from the Life sciences core facilities, Weizmann Institute of Science, Israel, for his guidance and technical support with the cryo-fluorescence imaging; Dr. Eyal Shimoni from the Electron microscopy unit, Department of research support, Weizmann Institute of Sciences, Israel and Dr. Assaf Gal for their help with the cryo-SEM/EDS analysis; Dr. Boris Shklyar at the Bioimaging unit, Faculty of natural sciences, University of Haifa, Israel, for his guidance and technical support with the in-vivo imaging; Dr. Jeana Drake for her advice on data interpretation and proofreading and to Maayan Neder, Itay Kolsky and Federica Scucchia for their help with sample collection.

## Author contributions

GMK and TL conceptualize the study and designed the experiments. GMK performed the experiments, GMK and SL analyzed the data. All the authors interpreted the data, prepared the initial draft. All the authors revised the manuscript and approved the final version for publication.

The authors declare that they have no competing interests

All data needed to evaluate the conclusions in the paper are present in the paper and/or the Supplementary Materials

## Supplementary Materials

Video S4: In-vivo laser scanning confocal fluorescence microscopy time-lapse (6min) of contraction and expansion movements of the tissue near the septa in a primary polyp.

## Supplementary Materials

**Fig. S1.**
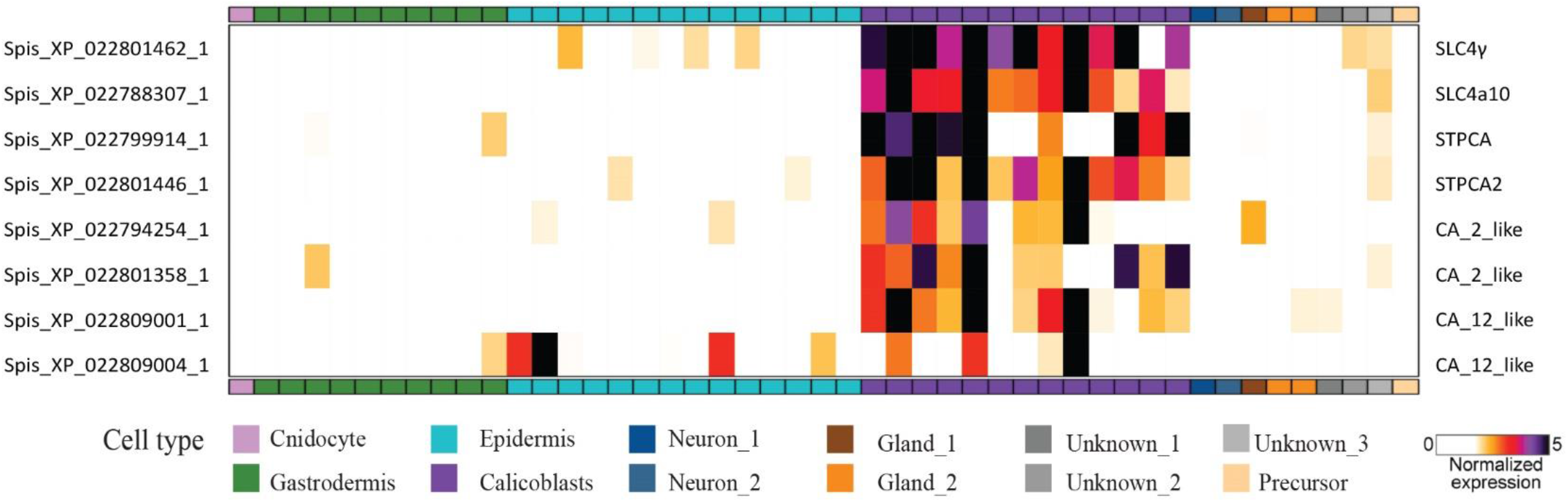
Expression heatmap of bicarbonate transporters and carbonic anhydrases genes across all cell types of *S. pistillata* primary polyp. All genes represented in this heatmap are highly enriched in the calicoblasts cells (represented by purple in the X-axis). (From *S. pistillata* primary polyp scRNAseq analysis *(80)*

**Fig. S2:**
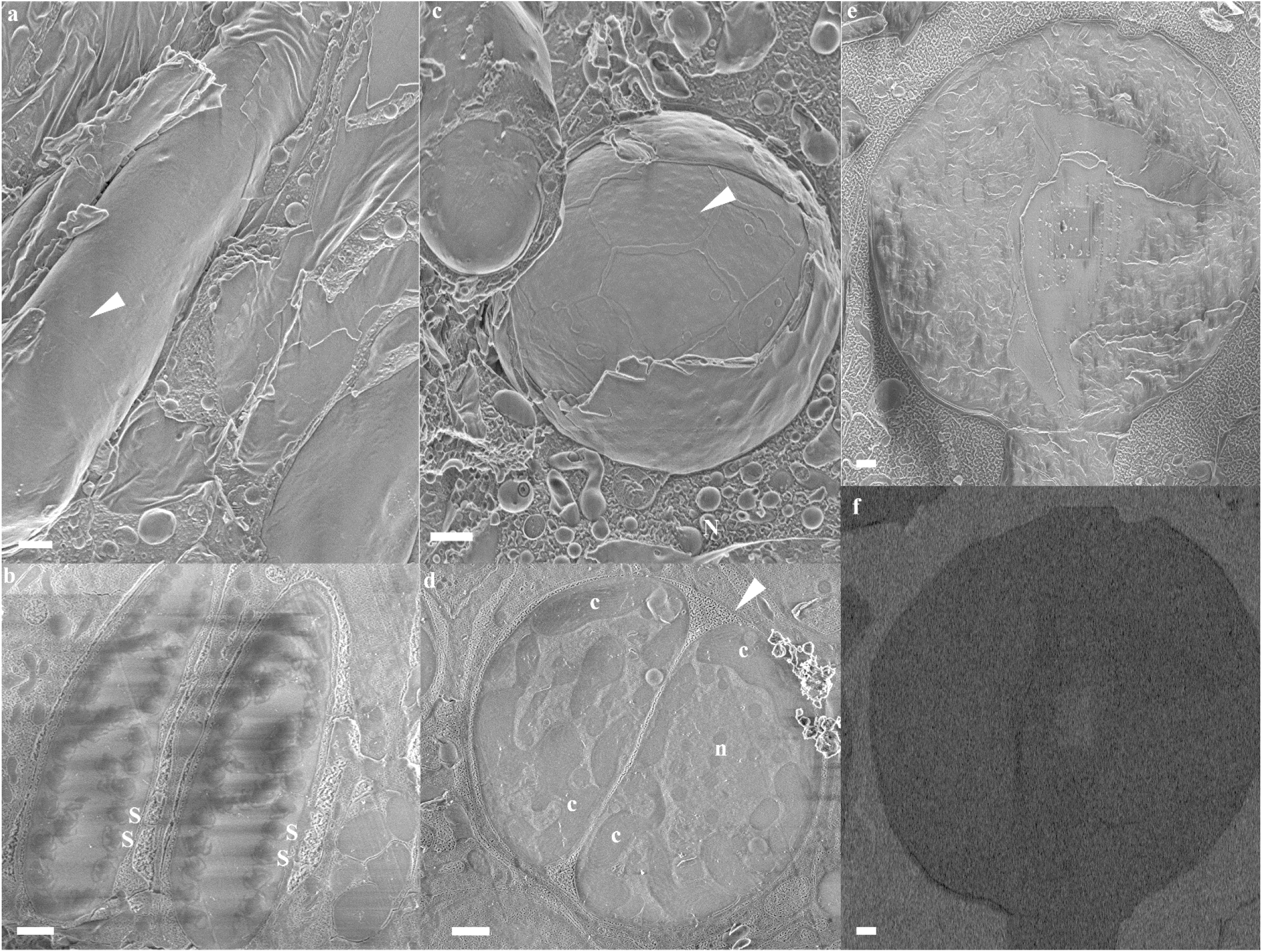
Cryo-SEM micrographs of oral tissue characteristic cells in primary polyps. (a) Nematocyte (white arrow head) imaged in a freeze fractured specimen (SE mode). (b) Two elliptical shaped nematocytes imaged in cryo-planed primary polyp (SE mode), each of them containing about 30 undischarged spiny tubules (stenotele), two representative stenotele are depicted in each cell. (c) Symbiotic algal cell imaged in a freeze-fractured primary polyp (SE mode) theca imprint on the cell outer surface are depicted with white arrowhead. (d) Dividing symbiotic algal cells within the coral host cell imaged in a cryo-planed primary polyp (SE mode). Host cell membrane depicted with white arrowhead. (e) A round shaped mucus cell imaged in a cryo-planed primary polp (SE mode) (f) the same image as (e) imaged in ESB mode, with bulk mucus cell content appear dark compared with its environment due to its organic content. All scale bars are 1µ m **S**- stenotele, **c**- chloroplasts, **n**-symbiont cell nucleolus.

**Fig. S3:**
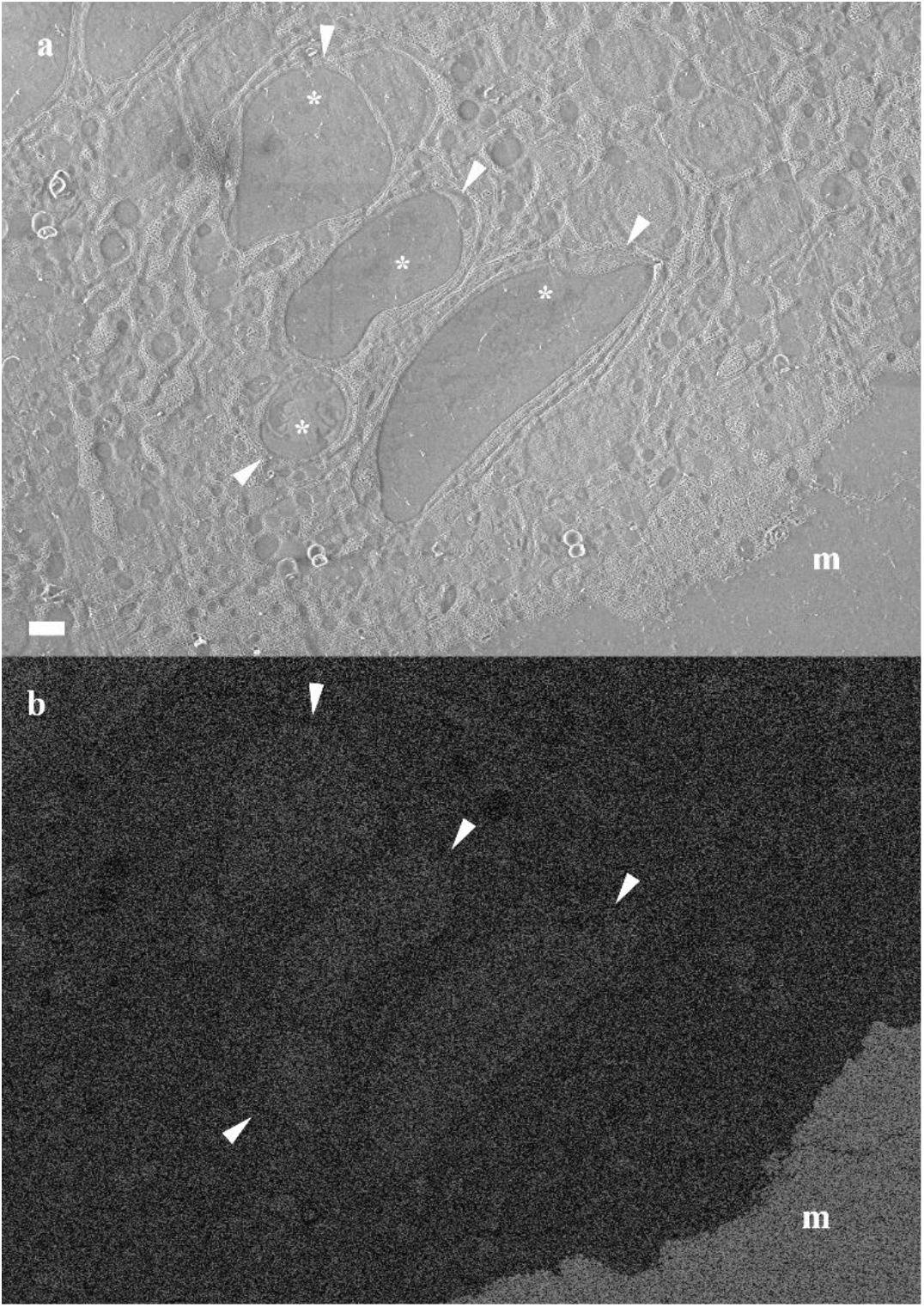
Cryo-SEM micrographs of nematocytes found in close proximity with the mineral. (a)SE mode. (b) ESB mode. Nematocytes are denoted with white arrowheads. Stenotele are marked with asterisk in image (a). **m**-mineral.

**Movie S4: In-vivo laser scanning confocal fluorescence microscopy time-lapse (6min) of contraction and expansion movements of the tissue near the septa in a primary polyp labeled with green fluorescent beads of 1 µm diameter**. The fluorescence images are composed of three overlayed channels: Green- fluorescence 1 µm sized beads, Red- symbionts auto-fluorescence and Grey scale-transmitted laser scanning image. A pumping movement is documented, created by contraction and extension of the ECM layer along all forming septa.

